# Spatial histomorphometry reveals that local peripheral nerves modulate but are not required for skeletal adaptation to applied load in mice

**DOI:** 10.1101/2024.05.12.593761

**Authors:** Alec T Beeve, Mohamed G Hassan, Anna Li, Nicole Migotsky, Matthew J Silva, Erica L Scheller

**Affiliations:** Department of Biomedical Engineering, Washington University in St. Louis, St. Louis, MO, USA; Division of Bone & Mineral Diseases, Department of Internal Medicine, Washington University School of Medicine, St. Louis, MO, USA; Department of Orthopaedics, Washington University School of Medicine, St. Louis, MO, USA

## Abstract

Mechanical loading is required for bone health and results in skeletal adaptation to optimize strength. Local nerve axons, particularly within the periosteum, may respond to load-induced biomechanical and biochemical cues. However, their role in the bone anabolic response remains controversial. We hypothesized that spatial alignment of periosteal nerves with sites of load-induced bone formation would clarify this relationship. To achieve this, we developed RadialQuant, a custom tool for spatial histomorphometry. Tibiae of control and neurectomized (sciatic/femoral nerve cut) pan-neuronal Baf53b-tdTomato reporter mice were loaded for 5-days. Bone formation and periosteal nerve axon density were then quantified simultaneously in non-decalcified sections of the mid-diaphysis using RadialQuant. In control animals, anabolic loading induced maximal periosteal bone formation at the site of peak compression, as has been reported previously. Loading did not significantly change overall periosteal nerve density. However, a trending 28% increase in periosteal axons was noted at the site of peak compression in loaded limbs. Neurectomy depleted 88% of all periosteal axons, with near-total depletion on load-responsive surfaces. Neurectomy alone also caused *de novo* bone formation on the lateral aspect of the mid-diaphysis. However, neurectomy did not inhibit load-induced increases in periosteal bone area, mineralizing surface, or bone formation rate. Rather, neurectomy spatially redistributed load-induced bone formation towards the lateral tibial surface with a reduction in periosteal bone formation at the posterolateral apex (-63%) and enhancement at the lateral surface (+1360%). Altogether, this contributed to comparable load-induced changes in cortical bone area fraction (+4.4% in controls; +5.4% in neurectomized). Our results show that local skeletal innervation modulates but is not required for skeletal adaptation to applied load. This supports the continued use of loading and weight-bearing exercise as an effective strategy to increase bone mass, even in patients with peripheral nerve damage or dysfunction.

**LAY SUMMARY:** Mechanical loading is required for bone health and can increase new bone formation to optimize strength and reduce fractures. Increased bone formation with loading is primarily mediated by osteocytes, a specialized cell type that is embedded throughout the bone matrix. Local nerve axons, particularly within the periosteum, may also respond to load-induced biomechanical and biochemical cues and could contribute to new bone formation in response to load. However, their role in the bone anabolic response remains controversial. To address this, we studied the adaptation of bone to applied load in the presence of local periosteal nerves, and in settings where the nerves had been surgically removed. Our results show that local skeletal innervation modulates but is not required for skeletal adaptation to applied load. This supports the continued use of loading and weight-bearing exercise as an effective strategy to increase bone mass, even in patients with peripheral nerve damage or dysfunction.

## INTRODUCTION

The skeleton is directly innervated by sensory and sympathetic nerves ^(1–3)^. Local axons integrate bone with the central nervous system to facilitate the conscious interpretation of local sensation and pain, and the rapid modulation of vascular tone in concert with other tissues throughout the body ^(1,4,5)^. Nerves in bone are concentrated around the arterioles and within the periosteum, with some potential for distribution into the marrow space ^(2,6)^. By contrast, the vascular sinusoids and endosteum are relatively aneural ^(2,6)^. Recent work has also shown that periosteal innervation is not uniform and contains three unique types neuroskeletal niches that have been defined based on the absence of nerve axons (Type I) and differences in the density and orientation of the nerve fibers (Type II/III) ^(2)^. The plasticity of these niches and their functional roles in sensation, pain, and anabolism are not yet understood.

Nerves in bone also have potential to release neurotransmitters near bone cells to modulate skeletal metabolism ^(1)^. Assessment of diverse studies over the past 100-years reveals that local nerves in and around bone (specifically) are likely to have neutral or mild modulatory functions on bone health *in vivo*, rather than an absolute requirement that links bone health to nerve function ^(7)^. Clinically, when peripheral nerve damage or dysfunction extends to other body systems, this has been associated with increased skeletal fragility in contexts of diabetes, spinal cord injury, chemotherapy-induced osteoporosis, multiple sclerosis, and anorexia ^(8–13)^. This clinical correlation can be due to shared mechanisms of disease onset between the skeletal and nervous systems. For example, recent work has shown that the rapid suppression of bone formation in type 1 diabetes is independent of the onset of diabetic peripheral neuropathy, though both present around the same time ^(14)^. Bone changes with neuropathy can also occur secondary to changes in other tissues such as muscle. Beyond this, it remains possible that the loss or modification of local neuronal cues may directly impact bone cell function, warranting context-specific clarification *in vivo*.

The skeleton adapts to its environment during weight-bearing exercise to strengthen bone. Upon loading, mechanical stimuli are sensed by osteocytes, matrix-embedded bone cells, which then signal to bone-forming osteoblasts and -resorbing osteoclasts to increase bone mass ^(15)^. Recent studies have suggested that the nervous system may be an additional gatekeeper for the anabolic response to load ^(16,17)^. If true, this could pose a significant problem for patients with nerve damage and dysfunction as most regimens to promote bone accrual rely on load-bearing exercise or movements. This proposed hypothesis warrants careful examination to determine exactly when and if the local nerves in bone contribute to the loading response.

There have been five prior studies that used chemical or surgical methods to study the role of the local peripheral nerves in load-induced bone formation ^(18–22)^. Specifically, Sample *et al* showed that temporary blockage of nerve activity via brachial plexus anesthesia reduced periosteal mineralizing surface after a single bout of very high 3750 με compression in the rat ulna (18N load), with no effects at lower, more physiologic, strains ^(18)^. Beyond this, four studies of unilateral surgical denervation of the leg observed either neutral or enhanced anabolic response to applied load in the denervated limb ^(19–22)^. This aligns with work by Heffner *et al* showing that global neonatal sensory denervation either had no effect or increased the anabolic response to applied load in adult animals depending on the site examined ^(23)^ and studies showing that neither sympathetic nerves nor β-adrenergic signals were required for the anabolic response to load ^(19)^. Altogether, these studies relatively conclusively show that local nerves are not required for load-induced skeletal adaptation. However, an important limitation of the prior work is that none of these studies visualized and confirmed loss of innervation in bone or related local nerve density to regional mineral apposition.

To address this, we developed RadialQuant, a custom tool for spatial histomorphometry. We first combined this with a Baf53b-tdTom pan-neuronal reporter mouse to develop a surgical model for the denervation of the tibia. Subsequently, we used RadialQuant to quantify and align load-induced bone formation and periosteal nerve axon density simultaneously in non-decalcified sections of the mid-diaphysis in either intact or denervated limbs. Our results show that local skeletal innervation modulates but is not required for skeletal adaptation to applied load. This supports the continued use of loading and weight-bearing exercise as an effective strategy to increase bone mass, even in patients with peripheral nerve damage or dysfunction.

## Methods

### Animals and biomechanical loading

#### Animals

This work was approved by the animal use and care committee at Washington University (Saint Louis, MO, USA). Pan-neuronal reporter mice (Baf53b-tdTom) were generated by crossing Baf53b-Cre (JAX Strain #027826) and Ai9-flox/flox (JAX Strain #007909) mice obtained from Jackson Laboratories, both strains were provided on a C57BL/6J background. Mice were group housed up to 5 mice per cage on a 12-hour light/dark cycle and fed *ad libitum* (PicoLab 5053, LabDiet). Unilateral nerve transection was performed at 16-weeks of age for all experiments. There were no adverse events.

#### Sciatic and Femoral Nerve Transection

Surgeries were conducted under anesthesia with 2% isoflurane as described previously ^(24)^. Briefly, for sciatic nerve transection, the lateral surface of the thigh was shaved and cleaned, and an incision was made to expose the sciatic nerve. A ∼5-mm portion of the sciatic nerve was then removed. For combined sciatic and femoral nerve transection, sciatic nerve transection was completed first. After this the medial surface of the thigh was prepared and an incision was made to expose the femoral nerve. A ∼5-mm portion of the femoral nerve was removed as above. The skin was closed with surgical staples (Durect Corporation, Reflex 7). Animals received a single 1.0 mg/kg dose of extended-release buprenorphine (ZooPharm) one hour before surgery for post-operative analgesia.

#### Strain Gauge Analysis

The use of transgenic mice in loading experiments requires strain gauge analysis to determine the peak force necessary to produce a desired strain on the surface of the tibia. To achieve robust lamellar bone formation at the mid-diaphysis of young-adult C57Bl/6 mice, a peak compressive strain of at least -2200με (peak tensile strain of 1200με) is required ^(25,26)^. To determine the peak force required to induce this strain, a strain gauge (FLK-1-11-1LJC, Tokyo Measuring Instruments Lab) was glued to the surface of the mid-diaphysis at the site of peak tensile strain in 16-week-old male and female Baf53b-tdTom mice. The tibia was then secured at the knee and heel in a custom fixture of a material testing system (Instron Dynamite 8841) and the strain gauges were connected to a data collection system (National Instruments SCXI-1001). Forces were applied to the tibia at 2N intervals between 2 and 10N, and the generated tensile strain was measured by the strain gauge and collected in LabVIEW (National Instruments). The force required to produce a 1200με tensile strain was interpolated from the resulting stress-strain curve (7.3N and 8.4N for females and males, respectively; Supplemental Figure 1). A previous study noted that woven bone formation occurs in the tibia at 10 and 11N applied force in male mice ^(26)^. Thus, to ensure lamellar bone formation in all animals without inducing woven bone formation, female mice were loaded at 8.5N and males were loaded at 9.5N for all experiments herein. This resulted in estimated peak tensile strains of 1395 and 1355με, respectively.

#### Biomechanical loading of the tibia

Axial compression of the tibia in the right hindlimb was achieved under 2-3% isoflurane anesthesia. The tibia was secured at the knee and heel in a custom fixture of a material testing system (Instron Dynamite 8841).

Compressive loading with 60-cycles at 4 Hz was applied every day for 5-days ^(25,26)^. Female mice were loaded at 8.5N and males were loaded at 9.5N for all experiments. Animals received a single 1.0 mg/kg dose of standard buprenorphine (ZooPharm) after loading for analgesia.

### Histology and Immunostaining

#### Tissue Isolation and Embedding

Mice were perfused under ketamine/xylazine with 10 mL PBS, followed by 10 mL of 10% neutral buffered formalin through the left ventricle (NBF, Fisher Scientific 23-245684). The left and right tibias were isolated from all animals, only removing the skin, and leaving muscle intact. Tibias were post-fixed in 10% NBF overnight then washed for 3 x 30 minutes in diH_2_O. For experiments that did not involve calcein labeling (only), tibias were then decalcified in 14% EDTA (Sigma-Aldrich E5134), pH 7.4. Tissues were infiltrated with 30% sucrose solution for 24-48 hours prior to embedding in OCT mounting media (Fisher HealthCare 23-730-571).

#### Cryosectioning of Decalcified Bone

For decalcified specimens, 50-µm-thick transverse tibial sections were cut on a cryostat (Leica) and mounted on Colorfrost Plus glass slides (Fisher Scientific 12-550-18). Step-by-step instructions are available at protocols.io (DOI: dx.doi.org/10.17504/protocols.io.bqu2mwye).

#### Tape Transfer Cryosectioning of Mineralized Bone

For calcified tibiae, 5-µm-thick transverse sections were cut at the mid-diaphysis on a cryostat set to -30 to -35°C (Leica) using a modified tape transfer protocol based on ^(27)^. In brief, we dispensed a drop of UV-curable adhesive (Norland Adhesive 63) onto TruBond 380 Adhesive Slides (Electron Microscopy Sciences 63700B10) and allowed the drop to run down the width of the slide to create a thin coat. Then, we applied an adhesive tape window (Leica 39475214) to the face of the block using a cold hand roller. We cut the block at 5 µm using high profile disposable steel blades (Leica 3802110) at a 0-2° clearance angle so that the section came off with the adhesive tape window. Tape-attached sections were mounted to the prepared slides, the adhesive was cured using a desktop UV illuminator (VWR M-20E) for 1 minute, and tape was removed from the slide using forceps.

#### Immunostaining

Step-by-step immunostaining instructions are available at protocols.io (DOI: dx.doi.org/10.17504/protocols.io.bqu2mwye) ^(2,28)^. In brief, sections were allowed to dry at room temperature (RT) for 30-minutes, followed by washing with PBS to remove OCT. Slides were blocked with 10% donkey serum in PBS, followed by permeabilization with 0.3% Triton X-100 in PBS for 1-hour at RT. For experiments that did not involve calcein labeling (only), slides were incubated with primary antibodies for calcitonin gene-related peptide (CGRP, Bio-rad, 1720-9007) and tyrosine hydroxylase (TH, Abcam, UK, ab152) at 1:1000 dilution overnight at RT in a humidified chamber. Then, slides were washed with PBS and secondary antibodies (Jackson Immunoresearch, 705-606-147, Jackson Immunoresearch, 711-546-152) were applied at 1:500 dilution for 3-hours at room temperature in a humidified chamber. After washing with PBS, slides were incubated once more with DAPI (Sigma-Aldrich) for 5 minutes and washed again thoroughly. A coverslip was mounted with Fluoromount-G (Thermo Fisher Scientific 00-4958-02).

#### Imaging

All slides were imaged using a Nikon spinning disk confocal microscope with a 10x objective (µm/px = 0.650, step size = 2.5 µm, Number of steps = 21 or 3 for 50- and 5-µm sections, respectively). Slides were imaged in tiles and stitched in the Nikon software prior to analysis as detailed below.

### Confocal Image Segmentation and Analysis

Investigators were blinded to the treatment condition during image analysis. Detailed step-by-step protocols for generation of bone masks, calculation of bone parameters using calcein, axon tracing and quantification, labeled surface tracing and quantification, and visualization of traced axons using image overlays can be found at protocols.io (DOI: dx.doi.org/10.17504/protocols.io.bp2l62b1dgqe/v1 (Private link for reviewers: https://www.protocols.io/private/9BBEACEA070611EFBAB60A58A9FEAC02 to be removed before publication). Written instructions are also available in the supplemental methods file.

### RadialQuant

A custom-made tool was written in MATLAB to align all bone images from the same location along the length of the bone and to partition the periosteum, axon, and labeled surface masks generated above into radial segments. Radial analysis facilitates the quantification of each feature locally within each radial segment, around the surface of the bone. All code and documentation are available open source on Github (https://github.com/alecbeeve/RadialQuant) ^(29)^. Additional details can be found in the supplemental methods.

### Statistics

Statistical analyses were performed in GraphPad Prism. Specific tests are indicated in the figure legends. A p-value of less than 0.05 was considered statistically significant.

## RESULTS

### Anabolic loading of the tibia increases bone formation without significantly increasing periosteal nerve density

Previous work suggested that nerves sprout toward bone in response to neurotrophic cues released from osteoblasts after mechanical loading ^(16)^. Thus, we hypothesized that mechanical loading would increase total axon density within the highly innervated periosteum where osteoblasts reside. To test this, we applied a unilateral axial load to the right hindlimb of Baf53b-tdTom male and female pan-neuronal reporter mice for 5-days with a regimen previously shown to induce lamellar bone formation ^(25,26)^ (Fig.1A, Day 1-5). Forces were determined based on strain gauging (see methods and Supplemental Fig.1). Calcein was administered at Day 4 and Day 9 after the first bout of loading with analysis of periosteal bone formation and axon density on Day 11 (Fig.1A). This was measured simultaneously around the entire circumference of the tibial mid-diaphysis in mineralized frozen sections (Fig.1B,C). In females, applied load caused a 4.6-fold increase in periosteal mineralizing surface (MS/BS) and a 24-fold increase in bone formation rate (BFR) compared to non-loaded limbs (Fig.1D,E). In males, the effect of applied load was less pronounced with no change in MS/BS and a 3.5-fold increase in BFR (Fig.1H). Though the loading paradigm brought about significant changes in bone formation, we did not observe a substantial difference in periosteal Baf53b+ nerve density after mechanical loading in either females (+15%, p = 0.321; Fig.1F) or males (+4%, p = 0.705; Fig.1I). Total axon length was similarly unchanged in both sexes (Supplemental Fig.2A,B).

**Figure 1.**
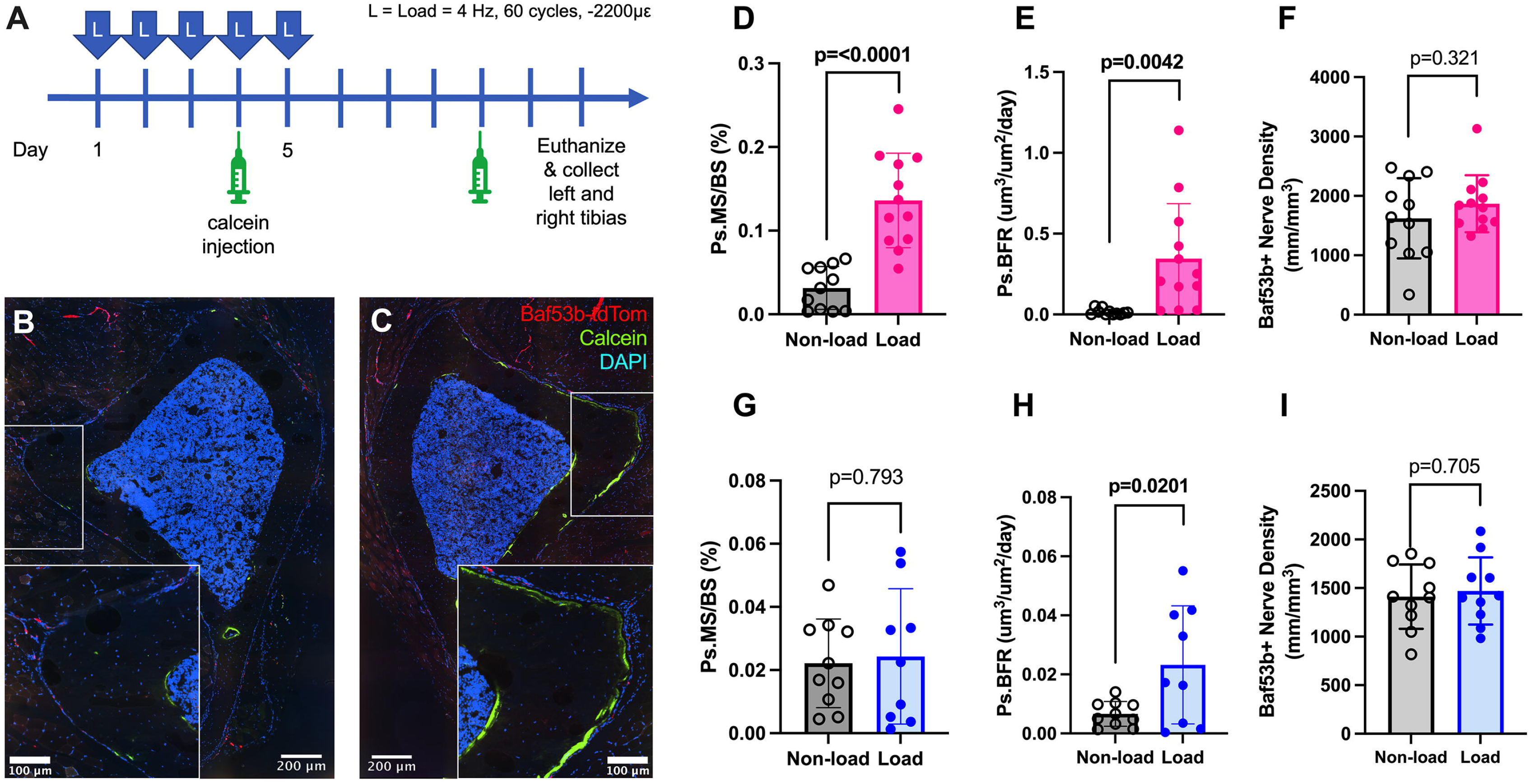
Axial compressive loading of the tibia increases bone formation but does not increase total periosteal nerve density 6-days after the last bout of loading. **(A)** Experimental timeline. **(B**,**C)** Representative confocal micrographs through a 5-um-thick transverse cross-section of a non-loaded **(B)** and loaded **(C)** tibia mid-diaphysis with DAPI, calcein labeling new bone formation, and tdTomato expression driven by Baf53b-Cre (nerves). Quantification of female **(D-F)** and male **(G-I)** periosteal mineralizing surface **(D**,**G)**, bone formation rate **(E**,**H)**, and Baf53b+ nerve density **(F**,**I)** as measured simultaneously around the entire circumference of the tibial mid-diaphysis. Data points represent individual animals with mean bar graph and standard deviation error bars; n = 9-12; unpaired t-test; bold p<0.05.

### Spatial distribution of periosteal nerves varies around the perimeter of the tibial mid-diaphysis and may change subtly with mechanical loading

Though the overall density of periosteal axons did not change significantly after loading, we hypothesized that periosteal axons may reposition towards actively mineralizing bone surfaces in response to applied load. To test this, we developed a novel spatial analysis tool called RadialQuant, which registers a set of 2D bone images and quantifies features of interest (*e*.*g*. nerves, mineral apposition) by segmenting the image into radial bins as it moves around a circle from 0 to 360° (Fig.2A). By evaluating only periosteal features with this approach, each radial bin corresponds to a specific region of the periosteal bone surface (Fig.2B). We sought to assess whether periosteal axon density varies around the surface of the mid-diaphysis, if the spatial distribution of axons is changed after loading, and whether variation in periosteal axon density aligns with patterns of mineralization induced by mechanical loading. As males had a modest biological response to loading, we focused RadialQuant analysis here on females only.

**Figure 2.**
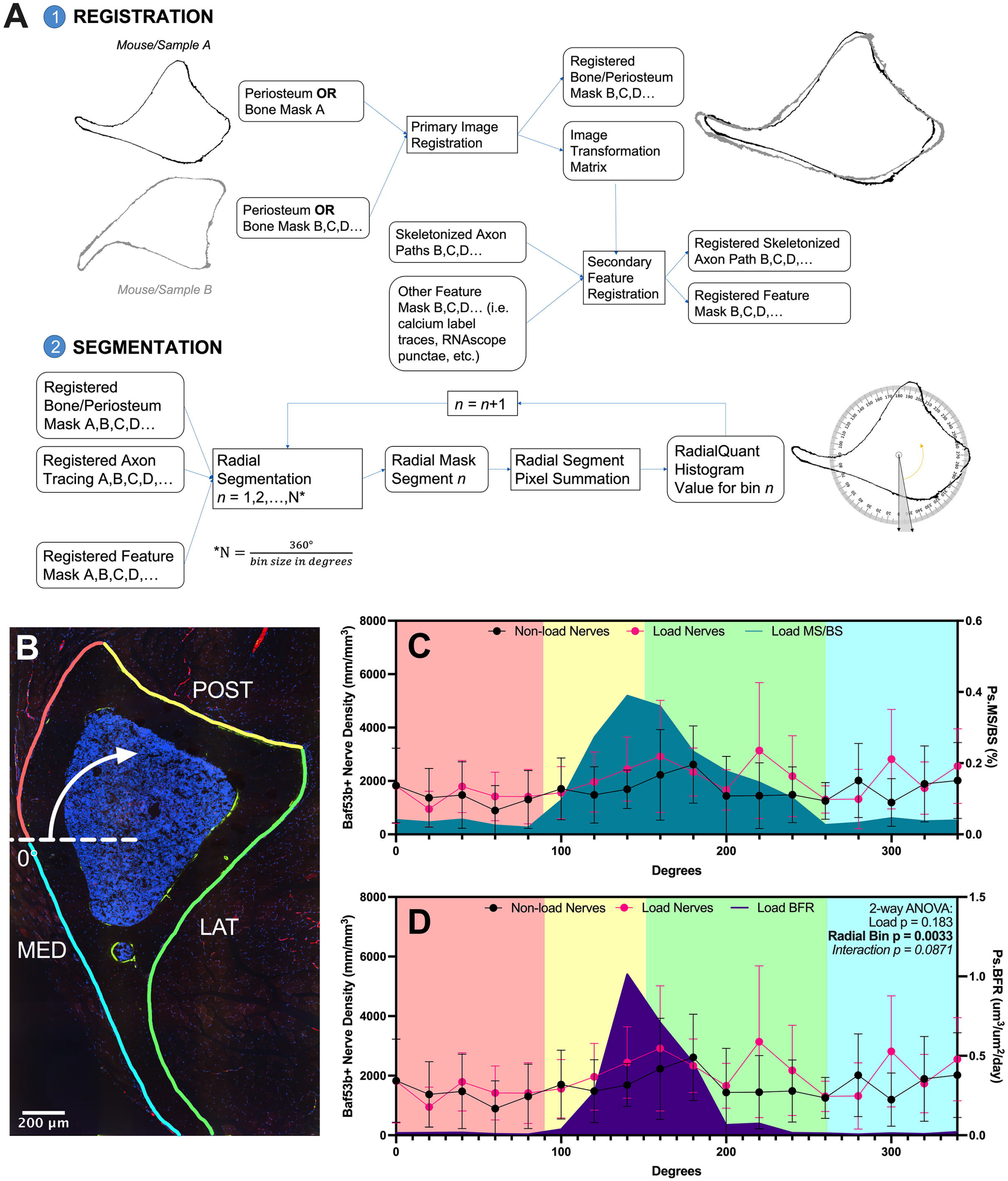
RadialQuant reveals a trending shift in the spatial distribution of local periosteal nerves in response to loading. Data shown for only for females, as they had high, consistent biological responses to loading. **(A)** RadialQuant workflow. **(B)** Representative confocal micrograph through a 5-um-thick transverse section of the tibia, labeled with four anatomical regions of the bone surface: postero-medial (red), posterior (yellow), lateral (green), antero-medial (blue). **(C**,**D)** RadialQuant histograms of mineralizing surface (C) and bone formation rate (D) overlaid on histograms for periosteal Baf53b+ nerve density. Shading represents the four regions of the bone surface listed in B. Data points represent the mean trendline with standard deviation error bars; n = 11-12; two-way ANOVA for nerve density; bold p<0.05, *italic* p<0.1. POST = posterior; MED = medial; LAT = lateral.

Periosteal axon density varied around the surface of the mid-diaphysis, with peaks at bins 140-180°, 220°, 300°, and 340°, corresponding to the posterolateral apex, the lateral surface, and portions of the anteromedial surface of the tibia (Figure 2C,D). Load-induced periosteal MS/BS and BFR also varied spatially around the surface of the mid-diaphysis, reaching a maximum at the site of peak compressive strain in the posterolateral apex (120° to 160°) (Fig.2C,D). A trending interaction effect indicated a shift in the spatial distribution of periosteal axons with mechanical loading (Baf53b+ nerve density, 2-way ANOVA, Load x Radial Bin p=0.087, Fig.2C,D). To determine if this aligned with sites of max skeletal adaptation, we evaluated periosteal axon density only at the posterolateral apex (120° to 160°). This identified a trending, but non-significant, 28% increase in localized periosteal axon density with loading (p=0.157; Supplemental Fig.2C,D).

### Denervation of the tibial periosteum requires both femoral and sciatic nerve transection

Though evidence of nerve sprouting was relatively limited after loading, we hypothesized that loading could also increase local signaling from periosteal nerves to enhance nearby bone formation. To test this hypothesis, we started by optimizing a protocol to surgically denervate the tibial periosteum at sites of maximum load-induced bone formation. Specifically, we tested sciatic nerve transection *vs* combined sciatic and femoral nerve transection in 4-month-old pan-neuronal Baf53b-tdTom reporter mice (Fig.3A). Periosteal nerves were quantified 7-days after nerve cut to allow for maximal degeneration ^(30)^. The contralateral limbs were used as intact controls.

**Figure 3.**
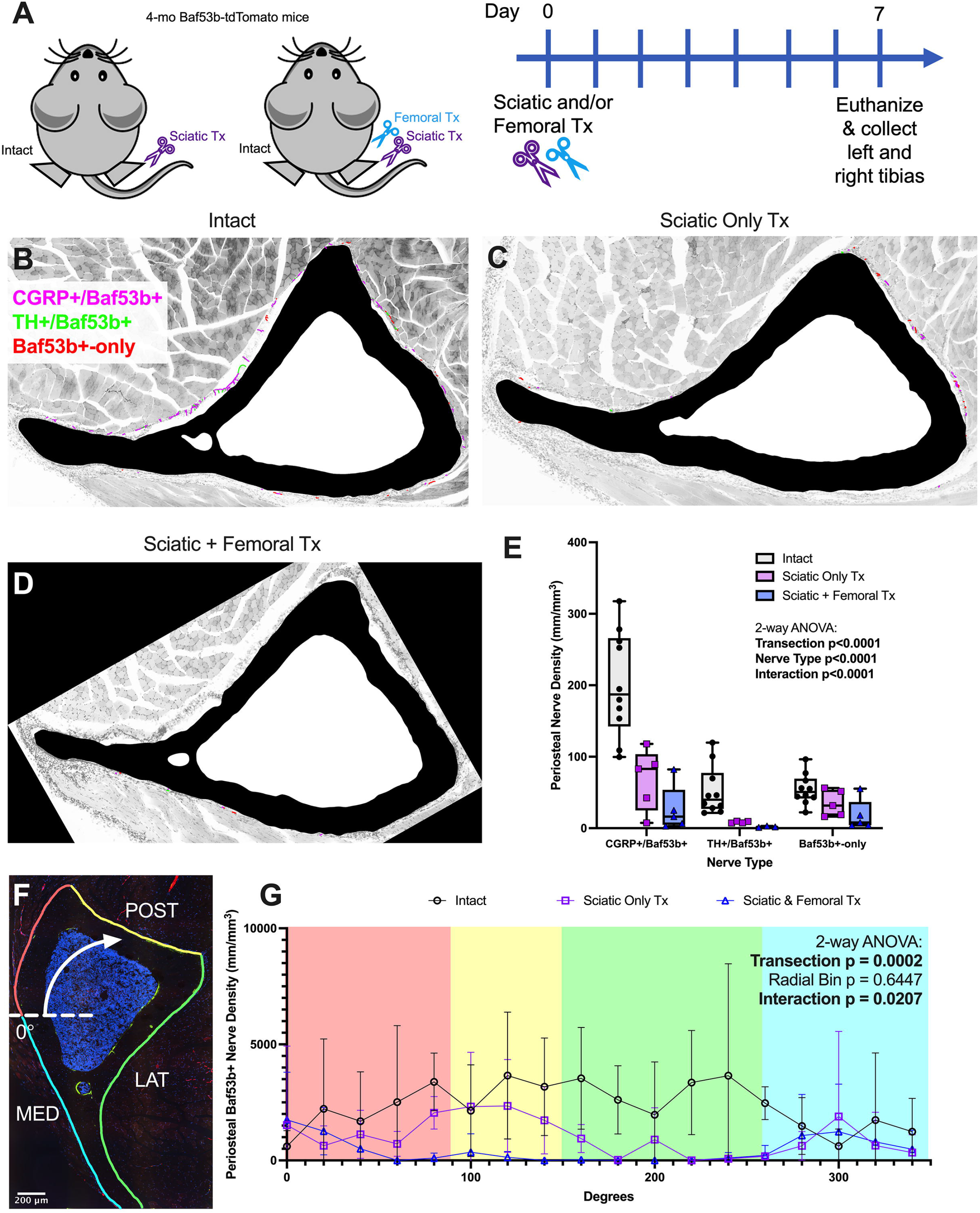
Denervation of the mouse tibial periosteum requires transection of both the femoral and sciatic nerves. **(A)** Experimental design and timeline. **(B-D)** CGRP+/Baf53b+ (pink), TH+/Baf53b+ (green), and non-peptidergic Baf53b+-only (red) nerve tracings overlaid on a cortical bone mask (black) and grayscale confocal image of surrounding fascia for representative intact (B), sciatic-only transection (C), and combined sciatic and femoral transection (D) experimental groups. **(E)** Periosteal axon density of CGRP+, TH+, and Baf53b+-only nerves. Data points represent individual animals; n=5-10; two-way ANOVA. Bold p<0.05 **(F)** Representative confocal micrograph through a 50-µm-thick transverse section of the tibia, labeled with four anatomical regions of the bone surface: postero-medial (red), posterior (yellow), lateral (green), antero-medial (blue). **(G)** RadialQuant histogram of all periosteal nerves (pan-neuronal Baf53b+) in intact, sciatic nerve cut, and sciatic/femoral nerve cut experimental groups. Shading represents the four regions of the bone surface listed in F. Data points represent the mean trendline with standard deviation error bars; n=5-10. Tx = transection; POST = posterior; MED = medial; LAT = lateral.

Sciatic nerve transection decreased periosteal innervation of the tibial mid-diaphysis by 67% (Fig.3B-E). When segmented based on immunostaining, this included 66% of sensory CGRP+ axons, 87% of sympathetic TH+ axons, and 36% of residual Baf53b+-only axons (Fig.3B-E). RadialQuant revealed that conserved axons after sciatic cut were primarily located along the posterior surface of the tibia with some extension onto the directly adjacent medial and lateral surfaces (Fig.3F,G). This residual innervation overlapped with the site of peak bone formation in response to applied load (Fig.2C,D). By contrast, combined sciatic and femoral nerve transection depleted 88% of total periosteal axons, including 87% of CGRP+ sensory fibers, 98% of TH+ sympathetics, and 67% of Baf53b+-only axons in the tibial mid-diaphysis (Fig.3B-E). This included complete depletion of periosteal nerves at the site of maximal load-induced bone formation (120° to 160°, Fig.3F,G). Conserved axons in the combined sciatic and femoral nerve transection group were localized to a limited region of the medial surface only, making this suitable for our proposed study (Fig.3G).

### Tibial denervation does not inhibit the anabolic response to load but spatially redistributes bone formation towards the lateral surface

To determine if local nerve-to-bone signaling is required to mount an anabolic response after applied load, we denervated the tibia via sciatic and femoral neurectomy 1-week prior to loading in 16-week-old male and female Baf53b-tdTom mice (Fig.4A). Loading forces were determined based on strain gauging of intact and neurectomized mice (see methods and Supplemental Fig.1). Notably, neurectomy 1-week prior to strain gauging did not significantly impact the force-strain curves, indicating no acute effect on overall bone mechanical properties (Supplemental Fig.1). There were four experimental groups: intact, load; intact, non-load; denervated, load; and denervated, non-load (Fig.4A). A 5 mm length of each nerve was removed to prevent re-innervation of the tibia. Consistent with this, persistent periosteal denervation was maintained throughout the study (Supplemental Fig.3).

**Figure 4.**
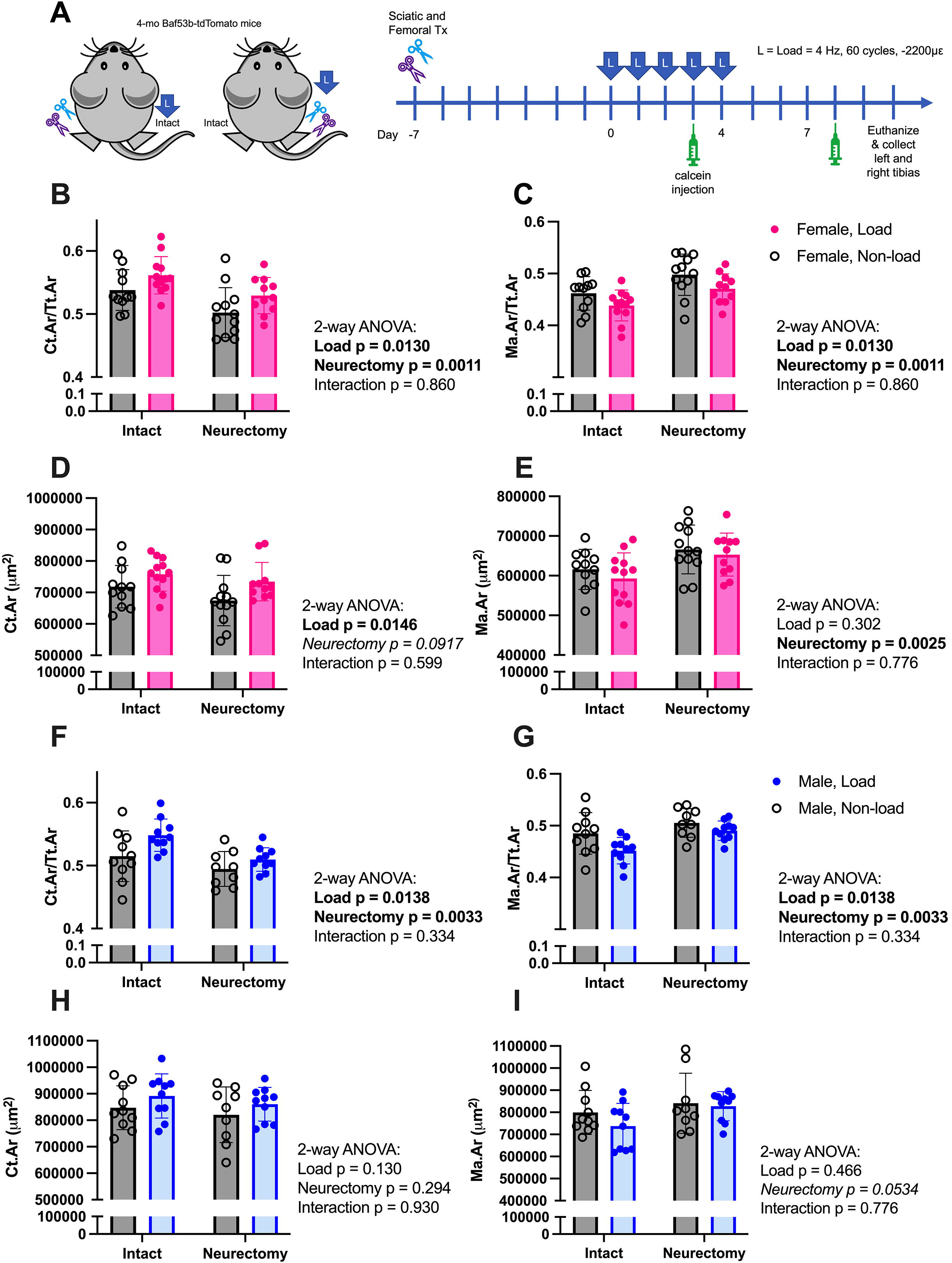
Local skeletal innervation is not required for load induced bone accrual. **(A)** Experimental design and timeline. **(B-E)** Female cortical bone data. **(F-I)** Male cortical bone data. Cortical area fraction **(B**,**F)**, marrow area fraction **(C**,**G)**, cortical area **(D**,**H)**, and marrow area **(E**,**I)** at the tibial mid-diaphysis. Data points represent individual animals with mean bar graph and standard deviation error bars; n = 9-12; two-way ANOVA; bold p<0.05, *italics* p<0.1. Tx = transection.

We started by analyzing bulk changes in cortical bone in non-decalcified sections of the tibial mid-diaphysis. In female mice, loading increased cortical bone area fraction in both intact and neurectomized limbs by 4.4% and 5.4%, respectively (Fig.4B), and reduced marrow area fraction in both intact and neurectomized limbs by 5.1% and 5.4%, respectively (Fig.4C). These effects were accompanied by load-driven increases in cortical bone area (Fig.4D) and a neurectomy-driven increase in marrow area (Fig.4E). In male mice, loading increased cortical bone area fraction by 6.4% and 3.0% in intact and neurectomized limbs, respectively (Fig.4F), and reduced marrow area fraction by 6.9% and 3.0% in intact and neurectomized limbs, respectively (Fig.4G). These effects were accompanied by a trending load-driven increase in bone area (Fig.4H) and a trending neurectomy-driven increase in marrow area (Fig.4I) that mirrored results in females. In both male and female mice, total area remained unchanged by load and neurectomy (Supplemental Fig.4). In summary, when considering bulk changes in cortical bone, load-induced increases in cortical bone mass were present in both males and females, regardless of neurectomy and despite neurectomy-induced increases in marrow area.

To study these changes in more detail, we performed dynamic histomorphometry using calcein double labels (Fig.5A). In females, loading significantly increased periosteal MS/BS and BFR, regardless of neurectomy (Fig.5B,C). In males, the effect of loading on dynamic histomorphometric indices was not significant (Supplemental Fig.5). In females, though the total amount of load-induced bone formation remained relatively constant (Fig.4,5), RadialQuant revealed that neurectomy substantially modified the spatial distribution of load-induced increases in MS/BS and BFR (3-way ANOVA, Radial Bin x Neurectomy x Load p<0.0001. Fig.5D-F). In intact limbs, load-induced increases in periosteal MS/BS and BFR peaked at 120° to 160°, corresponding to the posterolateral apex of the tibia mid-diaphysis (Fig.5D-F, Fig.6A-C). However, in neurectomized limbs, load-induced increases in periosteal MS/BS and BFR peaked from 200° to 240°, corresponding to the lateral surface (Fig.5D-F, Fig.6D,E). This shift aligned with regions of new bone formation on the lateral surface that were independently induced by neurectomy alone (Fig.5B,C, Fig.6). In summary, tibial denervation did not prevent the anabolic response to load but instead spatially redistributed load-induced bone formation towards the lateral tibial surface with a reduction in periosteal bone formation rate at the posterolateral apex (-63%) and enhancement of skeletal adaptation at the lateral surface (+1360%) (Fig.6).

**Figure 5.**
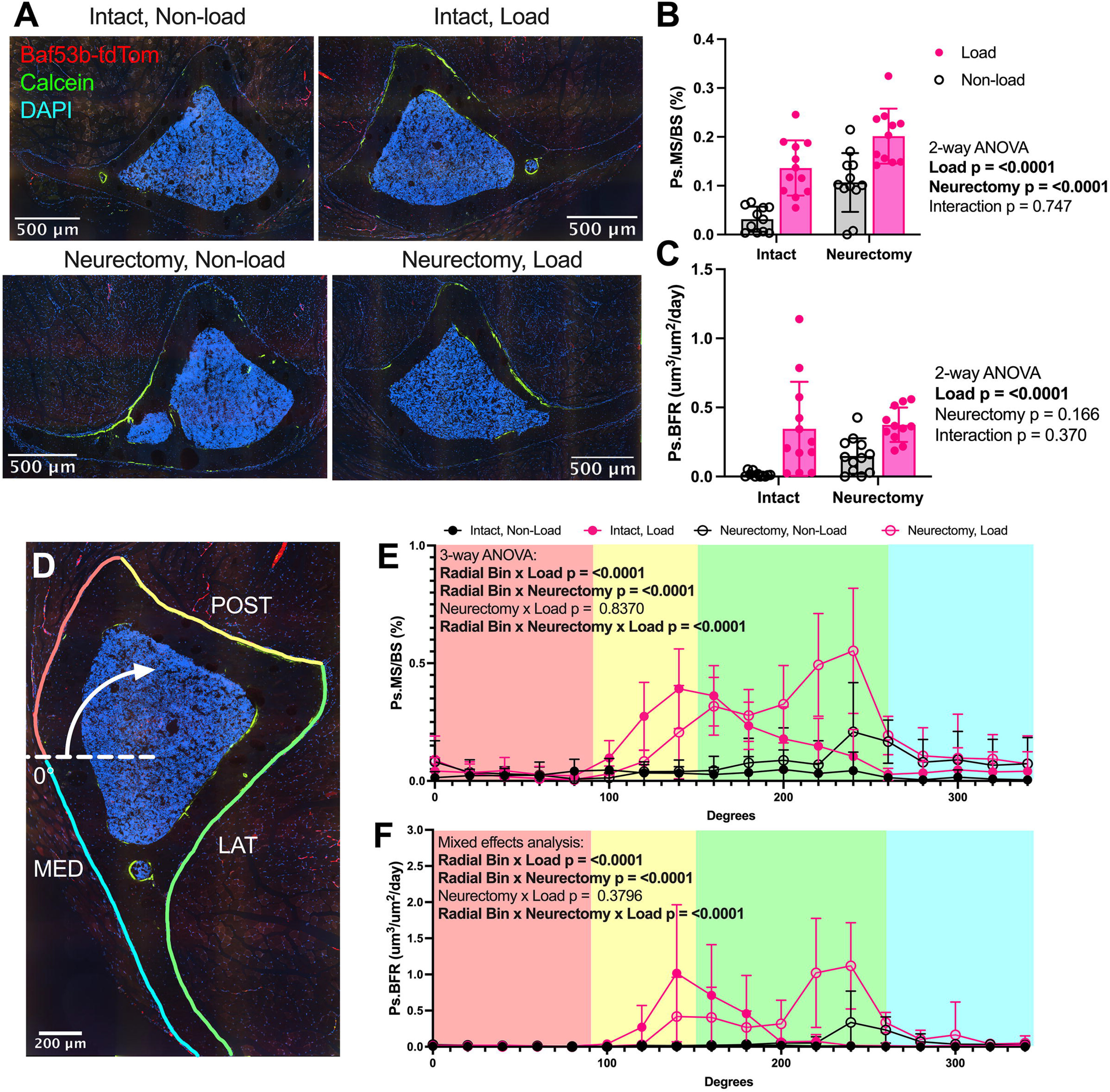
Denervation of the mouse tibia alters the spatial distribution of new bone formation in response to mechanical loading. **(A)** Representative confocal micrographs through a 5-µm-thick transverse cross-sections of the tibia mid-diaphysis of each experimental group. DAPI (blue), calcein labeling new bone formation (green), and tdTomato expression driven by Baf53b-Cre (red). **(B)** Periosteal mineralizing surface per bone surface (MS/BS). **(C)** Periosteal bone formation rate (BFR). Data points represent individual animals with mean bar graph and standard deviation error bars; n = 9-12; two-way ANOVA; bold p<0.05. **(D)** Representative confocal micrograph through a 5-µm-thick transverse section of the tibia, labeled with four anatomical regions of the bone surface: postero-medial (red), posterior (yellow), lateral (green), antero-medial (blue). **(E-F)** RadialQuant histograms of periosteal mineralizing surface (E) and periosteal bone formation rate (F). Shading in (E,F) represents the four regions of the bone surface listed in D. Data points represent the mean trendline with standard deviation error bars of each group; n = 11-12; 3-way ANOVA or mixed effects analysis with Tukey’s multiple comparison test; *p<0.050, ^†^p<0.10. POST = posterior; MED = medial; LAT = lateral.

**Figure 6.**
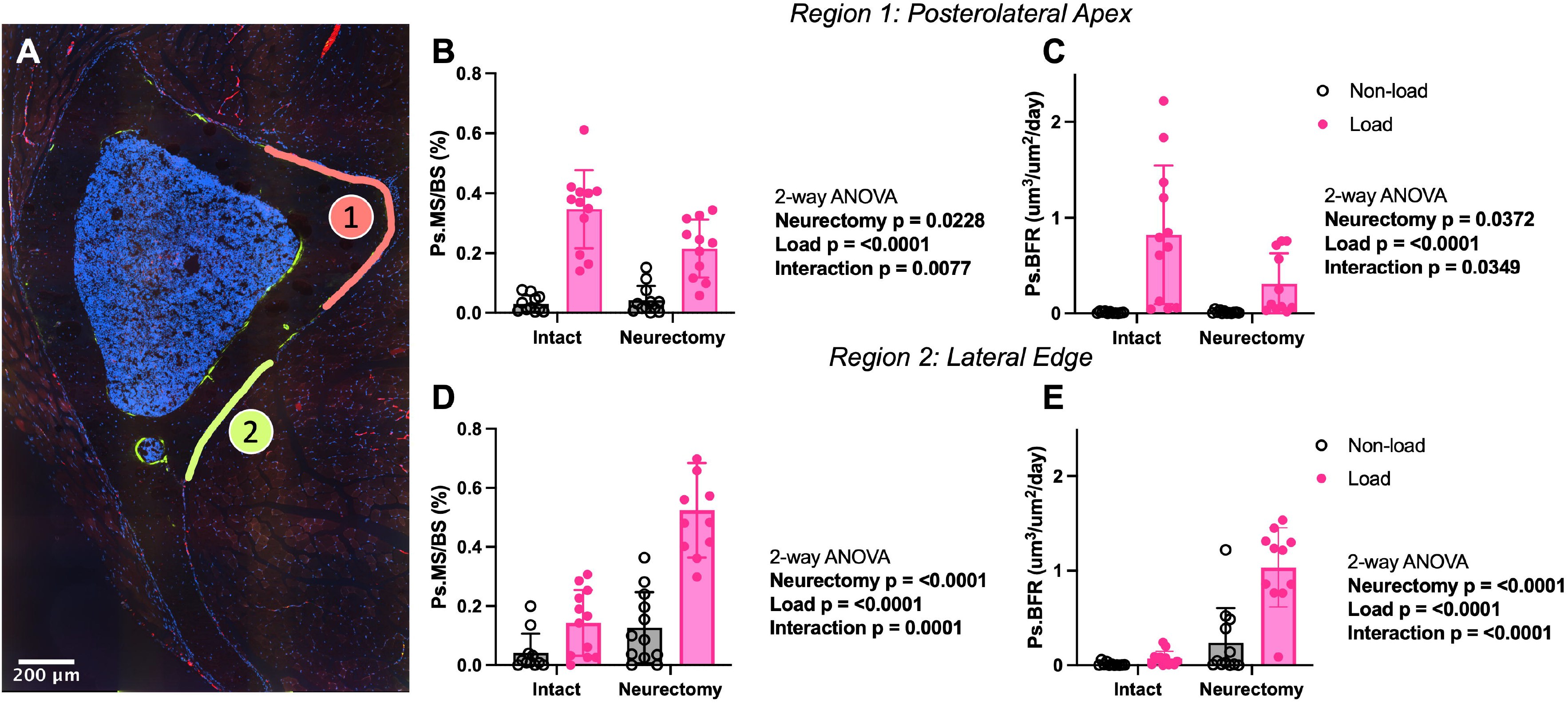
Neurectomy blunts load-induced bone formation at the posterolateral apex but enhances it on the lateral surface. **(A)** Representative confocal micrograph through a 5-µm-thick transverse section of the tibia, labeled with two anatomical regions of the bone surface: posterolateral apex (red, #1) and lateral edge (green, #2). Periosteal mineralizing surface per bone surface (MS/BS) **(B)** and periosteal bone formation rate (BFR) **(C)** at the posterolateral apex. Periosteal MS/BS **(D)** and periosteal BFR **(E)** at the lateral edge. Data points represent individual animals with mean bar graph and standard deviation error bars; n = 11-12; two-way ANOVA with Sidak’s multiple comparisons test; bold p<0.05.

## DISCUSSION

To assess whether periosteal axons were required for skeletal adaptation to applied load, we developed a model of tibial denervation. Consistent with clinical findings in humans ^(31,32)^, the mouse tibia was innervated by both the sciatic and femoral nerves. TH+ innervation of the tibial periosteum was 84% from sciatic, 12% from femoral, and only 4% from other sources. Consistent with work from the early 1900s, this suggests that sympathetic nerves in bone branch distally from the nerve trunks to innervate local arterioles rather than trafficking with larger arteries ^(22,33)^. Sensory peptidergic CGRP+ innervation of the tibia was 66% sciatic, 21% femoral, and 13% other. Lastly, Baf53b+ only nerves, which may represent a non-peptidergic sensory and/or non-adrenergic sympathetic subset, were 36% sciatic, 31% femoral, and 33% other. Axons from sciatic and femoral nerves distributed to distinct surfaces of the tibia, with the sciatic targeting the lateral and most of the medial surface and the femoral nerve contributing to periosteal innervation along the posterior edge. Additional sources of innervation that may explain the residual axons on the medial surface include the obturator nerve and smaller accessory branches from the lumbar region of the spine ^(34)^. Overall, this informs our foundational understanding of the innervation of the tibial periosteum and was sufficient to denervate the load-responsive surfaces of the tibia.

The hypothesis that nerve endings are the receptor of mechanical stress in bone was first put forward in 1937 by Hurrell, who also suggested a reflex arc governing bone growth and maintenance ^(35)^. The presence of mechanoreceptors was experimentally demonstrated in 2006 by members of the Rowe group, defining both the receptive field sizes (18 ± 14 mm^2^) and the mechanical response threshold (4.9 ± 4.3mN) of the individual periosteal neurons ^(36)^. Mechanoreceptive sensory nerves mediate the pain and sensation experienced with distortion of the periosteum due to excess bending and fracture of bone. Experiments to determine if these nerves were also required for the anabolic response to load were first initiated by Hert *et al* in 1971 ^(22)^. As in our study, they unilaterally resected both the sciatic and femoral nerves prior to quantifying the response of the tibia to applied load using x-ray and tetracycline-based dynamic histomorphometry, though in rabbits ^(22)^. Compressive stress was applied using implanted metal wires with a force of 1-2 kg at 1 Hz for 1-2 hours/day over 25-50 days^(22)^. Apposition of new bone formation with loading occurred regardless of denervation and to the same extent in denervated and intact limbs ^(22)^. Subsequent studies using sciatic denervation in mice with analysis at increased resolution have found either similar or enhanced responses to load after denervation based on the site and time point examined ^(19–21)^. This matches our findings in which the overall anabolic response to load was maintained regardless of denervation. Altogether, these results support the prevalent paradigm in which the osteocyte, and not the nerve ending, serves as the primary cell driving the anabolic response to applied load.

After mechanical loading, osteoblasts increase bone formation in regions of high strain, resulting in localized increases in bone mass. Though our data and others show that local nerves are not necessary for this response, it remains possible that changes in nerve function are sufficient to modulate the amount of bone formed. This could be through the hypersecretion of local neurotransmitters or through secondary changes in nervous system function throughout the body ^(7)^. Nerve growth factor (NGF) is one of the most highly and consistently upregulated genes in bone after compressive loading of the limbs ^(16,37,38)^. As most experimental loading regimens use high strain that can be painful, NGF secretion may coordinate a localized pain response with nerve sprouting to limit damage to the skeleton by alerting the host. In line with this, we observed a trending increase in nerve density at the site of peak compressive strain in females (+28%, p=0.157). In addition, RadialQuant of periosteal axon density showed a trending interaction effect of load and radial bin (2-way ANOVA, p=0.087), indicating possible spatial redistribution or mild sprouting of axons following mechanical loading. In addition to effects on nerves, NGF can activate TrkA receptors on endothelial cells and potentially also osteolineage cells to regulate *do novo* vasculogenesis and bone formation directly ^(7,39)^. Consistent with this, recent work found that when the high-affinity NGF receptor TrkA was blocked globally throughout the body, this prevented 41% of the load-induced increase in BFR, 16% of the load-induced increase in mineralizing osteoblasts, and 31% of load-induced increase in osteoblast activity ^(16)^. A global role for prostaglandin signaling in sensory nerves in the anabolic response to load has also been reported ^(17)^. Overall, when isolating the interactions between nerves and bone it remains important to consider the broad distribution of the nervous system throughout the body, the potential for secretion of neurotransmitters by non-neural cells, and the fact that most nerve-derived signals are also carried throughout the circulation ^(1,7)^.

The role of local nerves as modulators of the anabolic response rather than gatekeepers is also supported by our study, as new bone formation shifted from the posterolateral apex to the lateral tibial surface after denervation. Why? We observed an increase in BFR along the lateral edge of the tibia caused by neurectomy alone, indicating that sciatic and femoral neurectomy increases actively mineralizing osteoblasts after 1- to 2-weeks. One hypothesis is that transient increases in mineralization may be due to altered gait. Sciatic and femoral neurectomy causes dragging of the hindlimb and may result in non-habitual loading, which could explain the increased local apposition observed in the neurectomized limbs. Alternatively, degeneration of local axons may drive an acute increase in physiologic tissue function. This ‘regional acceleratory phenomenon’ secondary to injury or noxious stimulus was first described by Frost in 1983. This is not the first time this phenomenon has been documented. A transient increase in mineral apposition rate and BFR was noted in a previous study at 1-week after sciatic neurectomy in growing female rats ^(41)^. Though variable, similar increases in periosteal BFR on the lateral tibial surface were reported at 5- and 19-days after sciatic neurectomy in mice ^(20,21)^. Increased osteoblastic availability on the lateral surface secondary to neurectomy may have primed the subsequent responses to anabolic load in this region. It is important to note that responses to load in denervated limbs persist at levels at or above innervated bone long after this transient acceleratory phenomenon ends (even at 100-days after neurectomy), though it remains unknown if peak bone formation re-localizes to the apex ^(19,41)^. New RadialQuant spatial alignment and nerve tracing tools as reported here can help to answer these questions in the future.

As a final point, it is important to consider the clinical application of load-based exercise and movement to improve bone health in patients with neurologic disorders. Necessitating a neural relay to promote local bone formation vs relying on the mechanosensitive osteocyte network would be an inefficient evolutionary solution, particularly given the fragility of the peripheral nerve axons and potential for damage or loss. Consistent with this, load-based exercise is commonly used to support bone health in patients with peripheral neuropathy or nerve damage. This includes patients with diabetic peripheral neuropathy, cerebral palsy, spinal cord injury, chemotherapy-associated nerve damage, and other osteoporoses ^(42–47)^. Though it remains important to clearly elucidate situations in which the central and peripheral nervous systems actively modify the anabolic response ^(16,17)^, our results support the continued use of loading and weight-bearing exercise as an effective strategy to increase bone mass, even in patients with peripheral nerve damage or dysfunction.

## ACKNOWLDEGMENTS

This work was funded by grants from the National Institutes of Health (NIH) including the Skeletal Disorders Training Program T32-AR060719 and F31-AR081123 (ATB), R01-DK132073 (ELS), R21-DE032420 (ELS), R01-AR047867 (MJS) and R21-AR079052 (MJS). This work was also supported by the Musculoskeletal Research Center Cores at Washington University (P30-AR074992), microgrant funding from the Washington University Center of Cellular Imaging, and fellowship support from the Washington University Center of Regenerative Medicine (MGH). Special thanks to the Guilak and Meyer Labs for generously allowing use of their cryostats for some portions of the study.

## Notes

DISCLOSURE SUMMARY. The authors have no conflicts to disclose.

### Competing Interest Statement

The authors have declared no competing interest.

