## Supplemental Figures for "Spatial histomorphometry reveals that local peripheral nerves modulate but are not required for skeletal adaptation to applied load in mice"

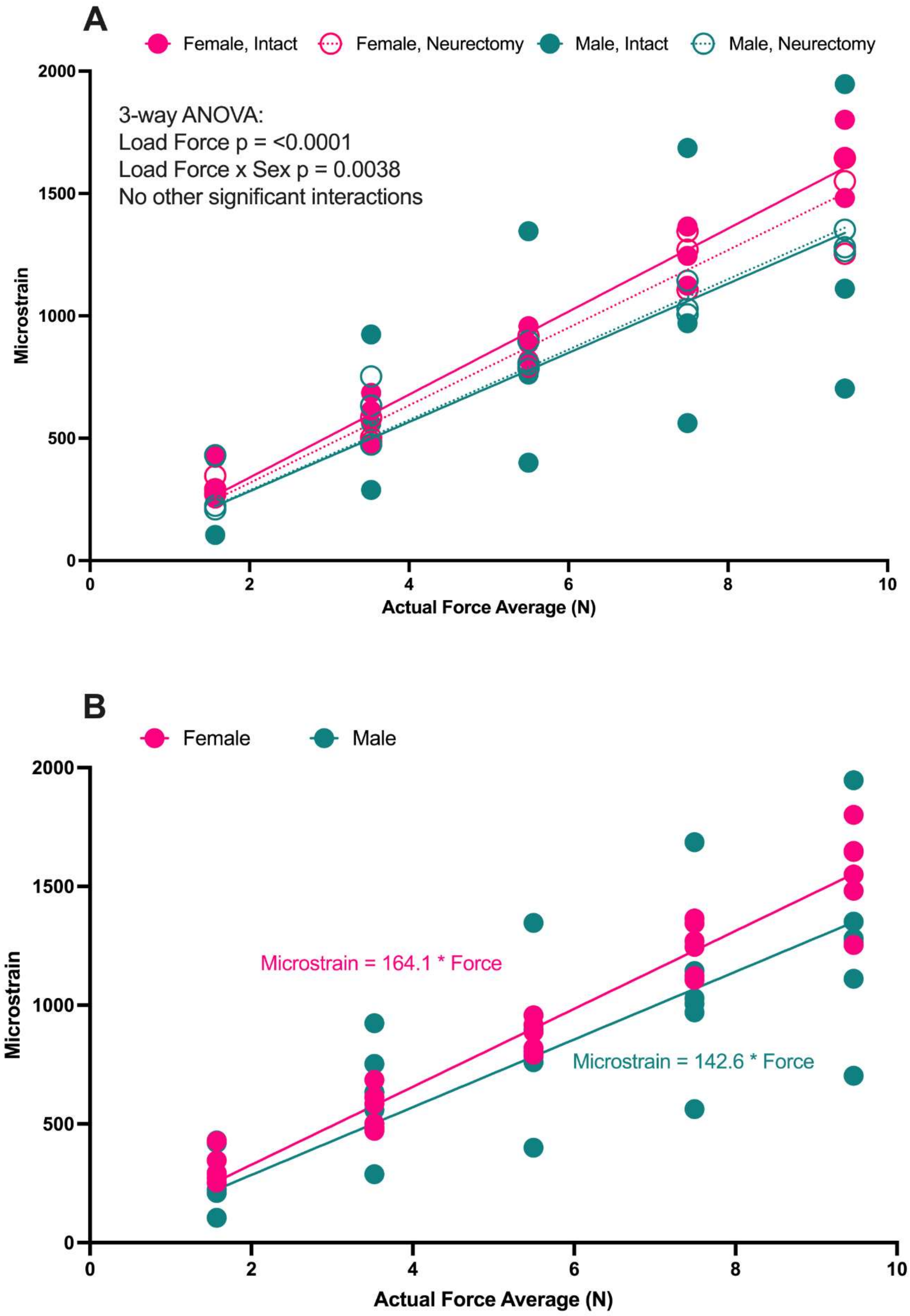


**Supplemental Figure 1. Strain gauge analysis. (A)** Stress-strain curve with intact and neurectomy groups separate. **(B)** Stress-strain curve with intact and neurectomy groups combined. Data points represent individual animals with mean trendline; n = 5-10; three-way ANOVA with Sidak's multiple comparisons test; bold p<0.05.

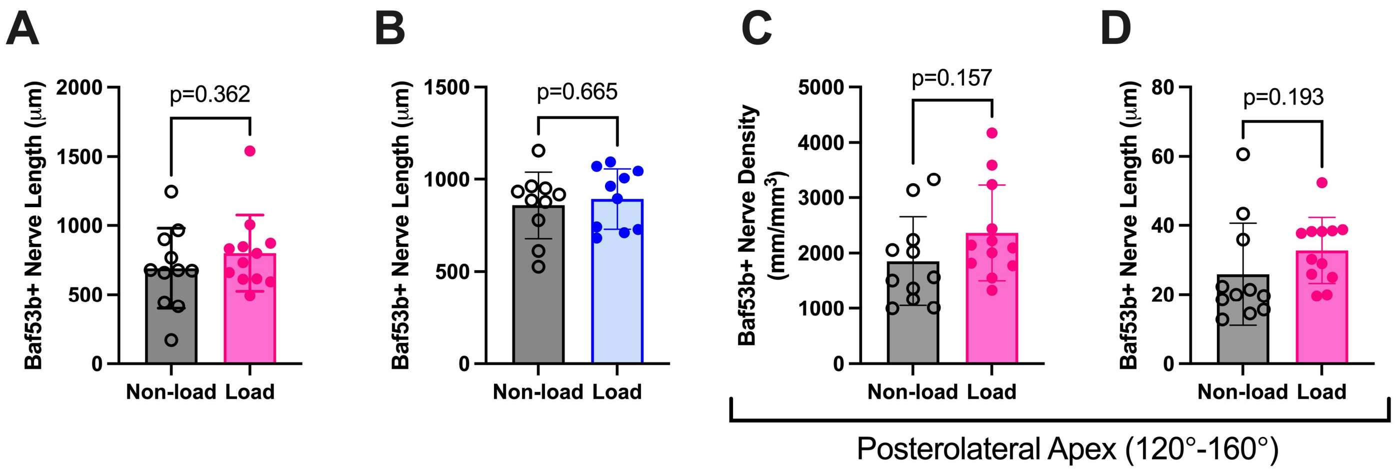


**Supplemental Figure 2. Pan-neuronal Baf53b+ nerve length within the whole periosteum and at the posterolateral apex with mechanical loading.** Female **(A)** and male **(B)** Baf53b+ nerve length in the total periosteum. Baf53b+ nerve density **(C)** and length **(D)** at the posterolateral apex in female mice. Data points represent individual animals with mean +/- standard deviation; n = 10-12; unpaired t-test; bold $p<0.05$; *italics* $p<0.1$.

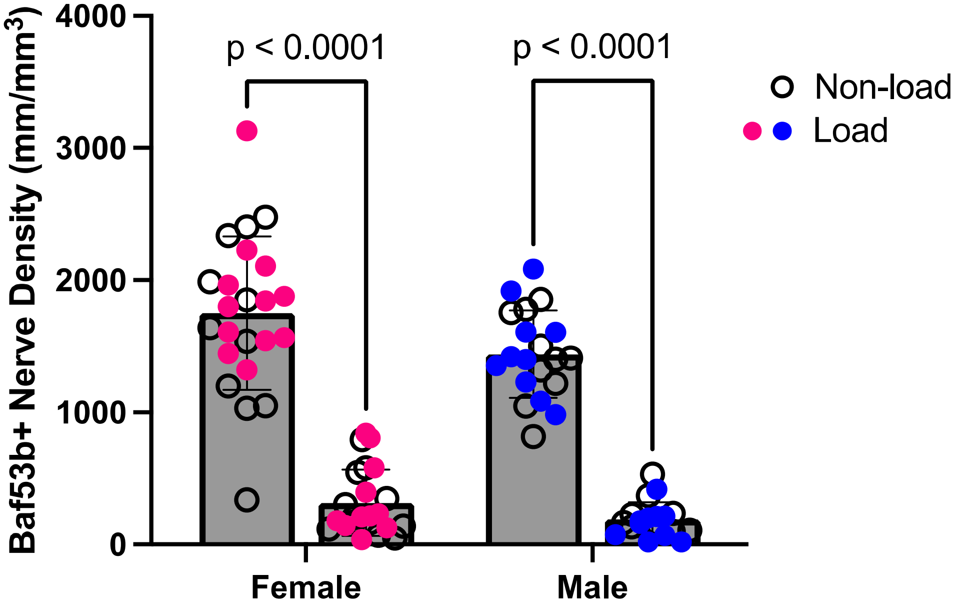


**Supplemental Figure 3. Total Baf53b+ periosteal nerve density at study endpoint 17-days post-neurectomy.** In order from left to right: female intact, female neurectomy, male intact, male neurectomy. Colored data points represent loaded limbs. Data points represent individual animals with mean +/- standard deviation; n = 10-12; 2-way ANOVA with Sidak's multiple comparison's test. P-values as indicated.

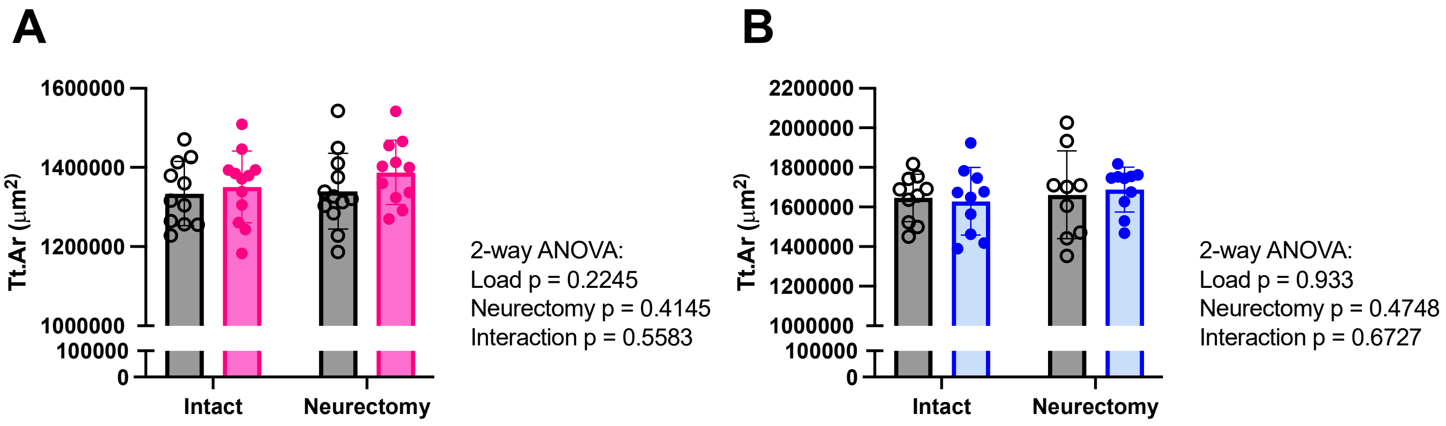


**Supplemental Figure 4. Quantification of total area with mechanical loading in intact and neurectomized limbs.** Female **(A)** and male **(B)** total area at the tibial mid-diaphysis. Data points represent individual animals with mean +/- standard deviation; n = 11-12; two-way ANOVA. P-values as indicated.

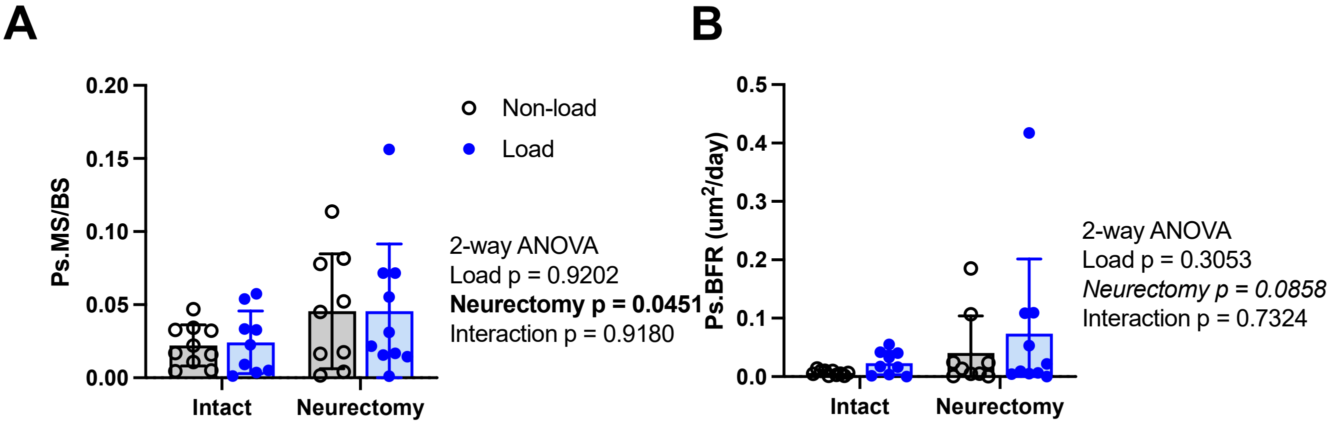


**Supplemental Figure 5. Male dynamic histomorphometry of the tibia with mechanical loading; intact and neurectomized limbs.** Periosteal mineralizing surface **(A)** and bone formation rate **(B)** at the tibial mid-diaphysis in intact and neurectomized limbs in male mice. Data points represent individual animals with mean +/- standard deviation; n = 9-10; two-way ANOVA; bold p<0.050; *italics* p<0.10.
