## Supplemental Methods for "Spatial histomorphometry reveals that local peripheral nerves modulate but are not required for skeletal adaptation to applied load in mice"

***Confocal Image Segmentation and Analysis***

Generation of bone masks. Generation of bone and periosteal masks was completed as described previously [^(1)^](https://sciwheel.com/work/citation?ids=10513470&pre=&suf=&sa=0&dbf=0), with some modifications. Specifically, bone masks were generated by thresholding max projections of the DAPI channel (blue) in FIJI [^(2)^](https://sciwheel.com/work/citation?ids=24178&pre=&suf=&sa=0&dbf=0), followed by manual correction to remove surrounding muscle and fascia. A median filter (radius=10) was then applied to the mask. Holes in the bone masks were filled using the FIJI binary operation Fill Holes. In cases where cortical bone completely enclosed the marrow cavity a small path was cleared from the cortex prior to performing the binary operation, and subsequently repaired. Finally, the mask was smoothed once again with a median filter (radius=30) and corrected manually as needed. Cortical area, total area, and marrow area were calculated from the bone mask using the Analyze Particles tool.

Generation of periosteal masks. The periosteum was defined as a thin, densely cellular layer on the external surface of the cortical bone. A threshold was applied to the DAPI channel once again to select the periosteum. Using previous masks, the bone and marrow were deleted from the periosteal mask. Masks were manually corrected to remove surrounding muscle and fascia. A Gaussian blur filter (radius=5) was applied to smooth the mask, followed by thresholding to produce a continuous layer surrounding the bone. After thresholding, the bone mask was utilized to remove areas of overlap with the periosteal mask. Lastly, holes in the periosteum were filled using the FIJI binary operation Fill Holes. Again, the mask was manually corrected as needed. The area of the periosteum was calculated using the Analyze Particles FIJI tool.

Bone surface mask and calculation. A bone surface mask was calculated by using the FIJI binary operation Fill Holes on the Bone Mask to fill the marrow cavity. Then, the FIJI binary operation Outline was used to create a bone surface mask. The total bone surface was calculated by utilizing the Analyze Particles FIJI tool for perimeter on the bone mask with marrow cavity filled.

Axon tracing and quantification. All but the periosteum was cleared from maximum projection images using the periosteal masks, and CGRP+, TH+ and Baf53b-tdTomato+ axons were traced additively in that order with the FIJI Simple Neurite Tracer plugin [^(3)^](https://sciwheel.com/work/citation?ids=4054583&pre=&suf=&sa=0&dbf=0) to obtain total axon length and single-pixel-width rendered path files for CGRP+ axons, CGRP+/TH+ axons, and CGRP+/TH+/Baf53b-tdTomato+ axons. With these data, we calculated axon densities for all axons, as well as CGRP+, TH+, and Baf53b+ only axon subtypes. The volume of the periosteum was calculated as the area multiplied by a section thickness of 50 or 5 µm. Volumetric density of fiber lengths was calculated as the total length of fibers traced divided by the corresponding volume for the region.

Labeled surface tracing and quantification. The FITC (green) channel of max projection images was utilized to trace and quantify single labeled surfaces (sLS), double labeled surfaces (dLS), and interlabel area (Ir.L.Ar). sLS were traced using the FIJI Simple Neurite Tracer plugin [^(3)^](https://sciwheel.com/work/citation?ids=4054583&pre=&suf=&sa=0&dbf=0) to obtain total single labeled surface and a single-pixel-width rendered file path of single labeled surfaces. dLS were traced and quantified similarly; only the midpoint of the inner label was traced. To quantify Ir.L.Ar, the midpoint of the outer label of all dLS was also traced. Then, double labels were enclosed on either end using a line segment tool in FIJI and subsequently filled using the FIJI binary operation Fill Holes. The FIJI Analyze Particles tool was used to quantify the total Ir.L.Ar.

Standardized parameters for dynamic histomorphometry parameters have been previously defined [^(4)^](https://sciwheel.com/work/citation?ids=1499222&pre=&suf=&sa=0&dbf=0). For this study, mineral apposition rate (MAR) was defined as the distance between midpoints of two consecutive labels, divided by the time between label injections. Mineralizing surface (MS/BS) was defined as the average labeled surface observed at the time of two label injections, normalized by the total bone surface (BS). Bone formation rate (BFR) was defined as the volume of mineralized bone formed per unit time, normalized by bone surface. Represented mathematically, these parameters were defined as:

$$MS/BS (\%)=\frac{dLS (\mu m)+\frac{sLS (\mu m)}{2}}{BS (\mu m)}$$

$$MAR=\frac{Ir.L.Th (\mu m)}{Ir.L.t (day)}$$

$$BFR (\mu m^{3}/\mu m^{2}/day )=MAR \left( \frac{\mu m}{day} \right)*\frac{MS}{BS} \left( \frac{\mu m}{\mu m} \right)$$

where

$$Ir.L.Th=interlabel thickness (\mu m)$$

$$Ir.L.t=interlabel time (day)$$

The computational approach described in this paper generated a new parameter, interlabel area (Ir.L.Ar), which is equivalent to the area between label midpoints at double labelled surfaces. Thus, we defined and calculated Ir.L.Th, as follows:

$$Ir.L.Ar \left( \mu m^{2} \right)=dLS \left( \mu m \right)*Ir.L.Th (\mu m)$$

$$Ir.L.Th (\mu m)=\frac{Ir.L.Ar (\mu m^{2})}{dLS (\mu m)}$$

Visualization of traced axons using image overlays. Single-pixel-width axon render path files were dilated four times using the FIJI binary operation Dilate, increasing the width of each traced axon to 9 pixels for the purposes of visualization. After this, bone and nerve masks were imported into Adobe Photoshop and overlaid on a grayscale max projection of the FITC and DAPI channels from the original image. This results in a visually accurate map of axon length, branching, orientation, and relative density that we have used to demonstrate the spatial orientation and positioning of the traced axons.

***RadialQuant***

Image registration. First, all masks (bone, periosteum, axons, labeled surfaces) produced in the study were transformed so that the cortical bone was oriented at the same angle. In brief, this was done by first selecting a universal “fixed” bone mask and orienting all other “moving” bone masks in the same direction as the “fixed” bone mask by creating three control points at the corners of the “fixed” and “moving” bone cross-sections to fit an affine geometric transformation matrix. The resulting matrix was then used to transform the remaining masks (periosteum, axons, labeled surfaces, etc.) into the same 2D orientation.

Radial segmentation and quantification. To partition the masks into radial segments, the bone or periosteum mask was utilized to determine the centroid of the analysis. Then, each mask was subsequently segmented into bins of 10-degrees by generating two rays with starting points at the centroid and end points at the edge of the image but with slopes 10-degrees apart. The number of pixels in the area between those two rays was recorded before the next set of rays was generated, with a 10-degree increase in slope from the former rays. This process continued until the final ray had a slope equivalent to 360-degrees. To reduce variability in the mid-diaphysis data with a 10-degree bin size, the bin size for this study was increased to 20-degrees computationally by summing raw data (periosteal area, axon length, sLS, dLS, Ir.Ar, etc) from two adjacent bins. Within each individual radial segment, periosteal axon density, MS, and BFR were calculated from raw data as described above.

**Associated References**

[1.    Lorenz MR, Brazill JM, Beeve AT, Shen I, Scheller EL. A neuroskeletal atlas: spatial mapping and contextualization of axon subtypes innervating the long bones of C3H and B6 mice. J. Bone Miner. Res. 2021 May;36(5):1012–25.](https://sciwheel.com/work/bibliography/10513470)

[2.    Schindelin J, Arganda-Carreras I, Frise E, Kaynig V, Longair M, Pietzsch T, Preibisch S, Rueden C, Saalfeld S, Schmid B, Tinevez J-Y, White DJ, Hartenstein V, Eliceiri K, Tomancak P, Cardona A. Fiji: an open-source platform for biological-image analysis. Nat. Methods. 2012 Jun 28;9(7):676–82.](https://sciwheel.com/work/bibliography/24178)

[3.    Longair MH, Baker DA, Armstrong JD. Simple Neurite Tracer: open source software for reconstruction, visualization and analysis of neuronal processes. Bioinformatics. 2011 Sep 1;27(17):2453–4.](https://sciwheel.com/work/bibliography/4054583)

[4.    Dempster DW, Compston JE, Drezner MK, Glorieux FH, Kanis JA, Malluche H, Meunier PJ, Ott SM, Recker RR, Parfitt AM. Standardized nomenclature, symbols, and units for bone histomorphometry: a 2012 update of the report of the ASBMR Histomorphometry Nomenclature Committee. J. Bone Miner. Res. 2013 Jan;28(1):2–17.](https://sciwheel.com/work/bibliography/1499222)
